# Human iPSC-derived prostate organoids with germline *BRCA2* mutation undergo tumorigenic transformations

**DOI:** 10.1101/2025.08.26.672478

**Authors:** Bipul R. Acharya, George Lawless, Pablo Avalos, Prince Anand, Yesai Fstkchyan, Shaughn Bell, Maria G. Otero, Samuel Guillemette, Zachary Myers, Michael Workman, William J. Catalona, Dan Theodorescu, Clive N. Svendsen

## Abstract

The lack of physiologically relevant *in vitro* prostate models has impeded studies of organ development and prostate tumorigenesis. We reprogrammed peripheral blood mononuclear cells (PBMCs) from individuals with and without pathogenic-germline *BRCA2* mutation (MUT_BRCA2, CON_BRCA2) into induced pluripotent stem cells (iPSCs), which showed no differences in morphology, proliferation, or pluripotency markers. Differentiation of MUT_BRCA2 iPSCs into prostate organoids (iPROS) using defined growth factors and signaling molecules resulted in disrupted morphology, impaired polarity, increased proliferation, and elevated prostate-specific antigen (PSA) secretion compared to CON_BRCA2 iPROS. Transcriptomic profiling revealed early prostate cancer (PCa) signatures. Upon exposure to dietary carcinogens, MUT_BRCA2 iPROS showed further PSA elevation, enhanced proliferation, AMACR upregulation, p63 reducetion are markers of aggressive PCa. *In vivo*, MUT_BRCA2 iPROS formed tumors in immunodeficient mice. This patient-derived iPROS-platform recapitulates human-prostate mopphology and function, models early tumorigenesis events, and provides a valuable tool for studying PCa biology and enabling personalized drug discovery.

**IN BRIEF:** In this study, we developed patients’ iPSC-derived prostate organoids (iPROS) with or without a pathogenic *BRCA2* germline mutation that display human-prostate like morphology and function. MUT_BRCA2 iPROS displayed disrupted morphology, early tumorigenic changes, and formed tumors in mice. Upon carcinogen exposure, they showed markers of aggressive prostate cancer. This platform models early prostate tumorigenesis and enables personalized studies of cancer initiation and therapeutic response.

## INTRODUCTION

Prostate cancer (PCa) is the most commonly diagnosed cancer in men globally, with over 1.2 million new cases and 350,000 deaths annually^1^. In prostate epithelium, genetic and epigenetic alterations, together with microenvironmental factors and oncogenic stress, often disrupt androgen receptor signaling, promoting PCa development and progression^2,3^. Among these alterations, germline mutations in DNA repair genes— particularly *BRCA2*—pose a significant challenge in managing PCa^4^. About 12% of men with metastatic PCa harbor such mutations, more than those with localized disease, and are less responsive to treatment^5^. Pathogenic-*BRCA2* mutation carriers have an 8.6-fold increased PCa risk, especially before age 65, and show poorer prognosis even with low-grade tumors. They also experience worse metastasis-free and PCa-specific survival following surgery or radiotherapy^6,7,8^. These tumors exhibit aggressive features, higher genomic instability, unique molecular profiles, and castration resistance, underscoring the need for translational models to guide therapy.

Although rodent models have provided insights into prostate development and cancer, significant anatomical and cellular differences with human prostate limit their translational value^9^. The human prostate is organized into distinct zones with a balanced basal-luminal cell ratio, whereas rodents have separate lobes and a luminal-dominant profile^10^. Additionally, access to fetal and adult prostate tissue is limited, and available cell lines are suboptimal for modeling human PCa^11^. Induced pluripotent stem cell (iPSC)-derived organoids offer an alternative by enabling the study of organogenesis, tumor initiation, and drug response in genetically defined, scalable systems^12^. Unlike tumor-derived organoids, which reflect late-stage disease and harbor pre-existing heterogeneity, iPSC-based models can recapitulate early oncogenic events by introducing genetic and non-genetic changes stepwise. They also allow generation of matched normal controls from the same genetic background, enhancing precision oncology applications^13^. Earlier models required rodent urogenital mesenchyme (UGM) for prostate specification from human pluripotent cells, limiting their preclinical utility^14,15^. Other embryonic stem cell-derived models without UGM don’t show any functional maturity^16^. In our study, we differentiated iPSCs generated from human peripheral blood mononuclear cells into prostate-like organoids (iPROS) using a rodent-UGM free, chemically defined system. These iPROS recapitulated key morphological, transcriptional, and functional features of the human prostate and further matured with vascularization upon xenotransplantation into immunodeficient mice^17^.

A major barrier in PCa research is modeling tumor initiation *in vitro*^18^. Controlled, human-relevant systems to define specific drivers of transformation are critical for risk prediction and therapeutic development^19^. To address this, we used dietary carcinogens to induce tumorigenesis in iPROS. PhIP, a heterocyclic amine found in cooked meat, and MNU, a potent DNA alkylating agent, are known rodent PCa inducers^20,21^. PhIP undergoes P450-mediated activation, forming DNA adducts that drive mutations and genomic instability^22^, while MNU introduces O6-methylguanine lesions that mispair during DNA replication, causing G-to-A transitions^23^. We exposed iPROS to these carcinogens to model tumorigenic molecular and morphological transitions^24^.

Our iPROS system closely mimics human prostate morphology and function, offering an ethical and accessible model for studying both organ and cancer development, identifying early biomarkers, and evaluating therapy responses. Importantly, it enables investigation into the mechanistic impact of *BRCA2* mutations on PCa onset and progression, advancing personalized medicine^25^.

## RESULTS

### Development of prostate organoids (iPROS) from patient iPSCs

To establish a robust model for prostate organoid differentiation, we began with three control iPSC lines derived from PBMCs in the Cedars Sinai iPSC core facility (**Table S1**). Prostate originates from urogenital sinus (UGS)—a caudal extension of the hindgut—formed from definitive endoderm (DE) during late embryogenesis (gestational weeks 10 to 12) and completes maturation at puberty^10,26^. Given the known role of signaling pathway modulation in embryonic-development, we designed a stepwise differentiation protocol from iPSC to DE, hindgut endoderm (HGE), and subsequently, iPROS (**Fig. 1A**). To efficiently induce DE, we treated hiPSC with CHIR99021 (a GSK-3β inhibitor), Activin A, and progressively increasing serum concentrations. Since UGS arises from hindgut, we directed DE toward hindgut lineage by activating WNT3A and FGF4 signaling^27,28^. FGF10 and WNT10B are essential during prostate development and branching morphogenesis, with FGF10 driving early bud formation and WNT10B potentially aiding in prostate specification^29^. After 48 hours of HGE induction, we supplemented culture with FGF10 and WNT10B while reducing FGF4 to promote prostate-specific specification over the next 48 hours, formed tiny-3D spheroids. 3D spheroids were them embedded in Matrigel and cultured them for 5 days with high levels of Dihydroxy testosterone (DHT), and andromedin factors FGF7, and FGF10, mimicking the urogenital mesenchyme signals that drive *in vivo* prostate budding and urogenital epithelial (UGE) development^26,30^. Additionally, we included SAG (a Sonic Hedgehog agonist) to support UGE differentiation into basal and luminal cells^31^. From day 15 onwards, until Hayflick limit achieved (∼14-16 weeks)^32^, we cultured them in a low-dose DHT medium alongside various growth factors, activators, and inhibitors detailed in the Methods. Media formulation was informed by prostate-development literature and adapted from previous studies^30,16,33,14,15^. After day 15 mechanical dissociation of large-organoid-mass (**Fig. S1A**), round-organoids grow singly for weeks, then form complex structures, merging into masses that periodically require mechanical separation. Prostate epithelial buds typically emerge from UGS and branch to form glandular ducts comprising luminal and basal layers. iPROS recapitulated such ductal structures at around 8-week (**Fig. 1B**). Immunohistochemistry and immunofluorescence (IF) confirmed expression of prostate-specific markers—Androgen Receptor (AR), NKX3.1, Prostate Specific Antigen (PSA), and lineage-specific markers: CK8-18 (luminal), p63 (basal), and Chromogranin A (neuroendocrine) (**Fig. 1C,D** and **S1B**). PSA, a hallmark luminal secretion product^34^, appeared in the media by week 6, increasing to ∼60 pg/mL by week 12, indicating organ-maturation (**Fig. 1E**). We next validated prostate-specific gene expression in iPROS via RT-qPCR (**Fig. 1F**). Compared to *GAPDH* (Avg. Cq = 18), target genes *AR*, *NKX3.1*, *p63*, *KLK3* (PSA), and *CK18* displayed Cq values of 20–35, reflecting moderate to high RNA abundance. To assess transcriptomic similarity between iPROS and native prostate, we performed mRNA-seq on iPROS and compared the data with 282 normal prostate samples from the GTEx Portal^35^. A panel of 30 highly expressed non-ribosomal genes showed expression patterns resembling those of adult prostate tissues (**Fig. 1G**), with 80% transcript overlap confirmed (**Fig. 1H**) (hypergeometric test, p-value = 4.445679e-10). Further comparisons using two independent adult prostate mRNA-seq datasets from GEO revealed ∼70% similarity, reinforcing the relevance of the model (**Fig. 1H**).

**Figure 1:**
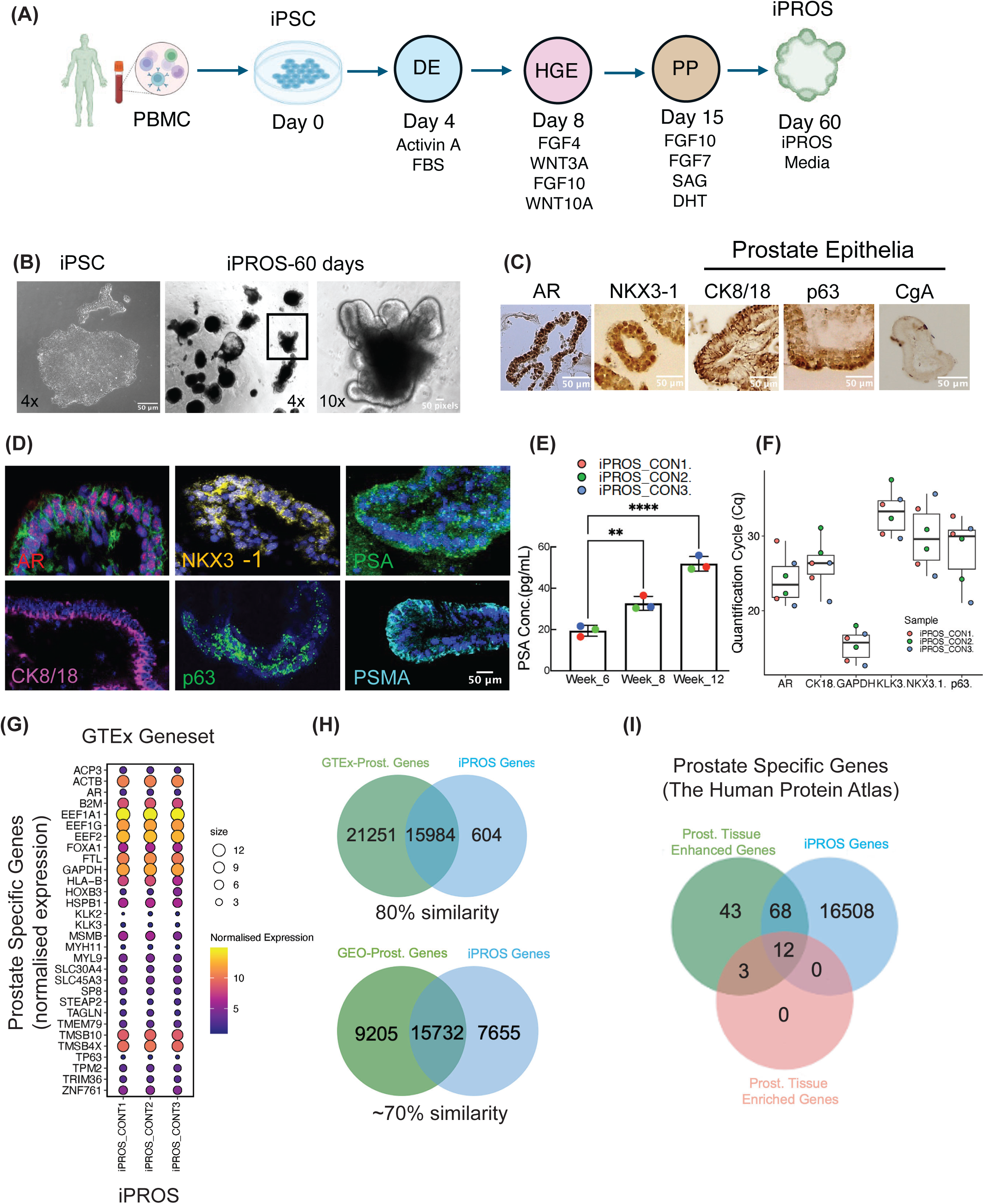
Differentiation and characterization of iPROS. (**A**) Prostate organoid differentiation-schema from iPSC_to_iPROS. (**B**) Representative images of iPSC and iPROS at day 60. (**C**) Chromogenic images showing AR, NKX3.1, CK8/18, p63, and CgA. (**D**) IF-images showing AR, NKX3.1, CK8/18, p63, PSA, and PSMA with DAPI. (**E**) Quantification of PSA release over weeks. (**F**) RT-qPCR Cq plot for prostate-specific genes. (**G**) Heatmap of 30 highly expressed prostate-specific genes from GTEx. (**H–I**) Venn diagrams showing gene overlaps of iPROS with GTEx, GEO, and Human Protein Atlas. Significance tested with ANOVA (**E**); **p < 0.01, and ****p < 0.0001.

Additionally, we queried the Human Protein Atlas and identified 126 genes showing >4-fold higher expression in prostate vs. other tissues (Prost. Tissue Enhanced Genes), with 15 “Prost. Tissue Enriched Genes” uniquely elevated in prostate as “tissue-specific genes”. Of these, 68 and 12 genes, respectively, were expressed in iPROS, with significant overlap (pValue = 1.115628e-06 and 2.322562e-10, respectively) (**Fig. 1I**). Lastly, we compared iPROS transcript profiles to cell-type specific mRNAs identified by total RNA-seq of adult prostate epithelium and stroma^36^. Heatmaps show normalized expression of luminal, basal, neuroendocrine, and stromal fibroblast markers in iPROS (**Fig. S1C**). These demonstrate that we developed a human prostate organoid model that mimics cell-specific morphology, gene expression, and organ function.

### iPROS with germline BRCA2 mutations manifest tumorigenic morphology and molecular signatures

Pathogenic-*BRCA2* germline variations are a known genetic risk factor for aggressive and metastatic PCa^5^. To assess their functional impact on iPROS morphology, gene expression, and PSA secretion, we generated three additional iPSC lines from PBMCs of three consenting PCa patients from Northwestern University (IRB#STU00018651-MOD0018) harboring pathogenic *BRCA2* germline mutations. **Fig. 2A** (**Fig. S2A** and **Table S1**) presents patient identification, mutation types, ISUP_Gleason-grade-group (GG) scores, and H&E-stained prostatectomy sections. Patients 1 and 2 share same 5946delT mutation in exon-10 but had differing GG scores: patient 1 (MUT1_BRCA4i; pT2) had GG2; patient 2 (MUT2_BRCA2A; T2) had GG5. Patient 3 (MUT1_BRCA3i) had a c.9513_9516 deletion in exon-11, GG5, and a pathology stage of T3bN1M1, with lymph node metastasis and histology revealing densely packed glandular lumens with fibrotic stroma. All mutations remained present in iPSCs and subsequent iPROS (**Fig. 2B**).

**Figure 2:**
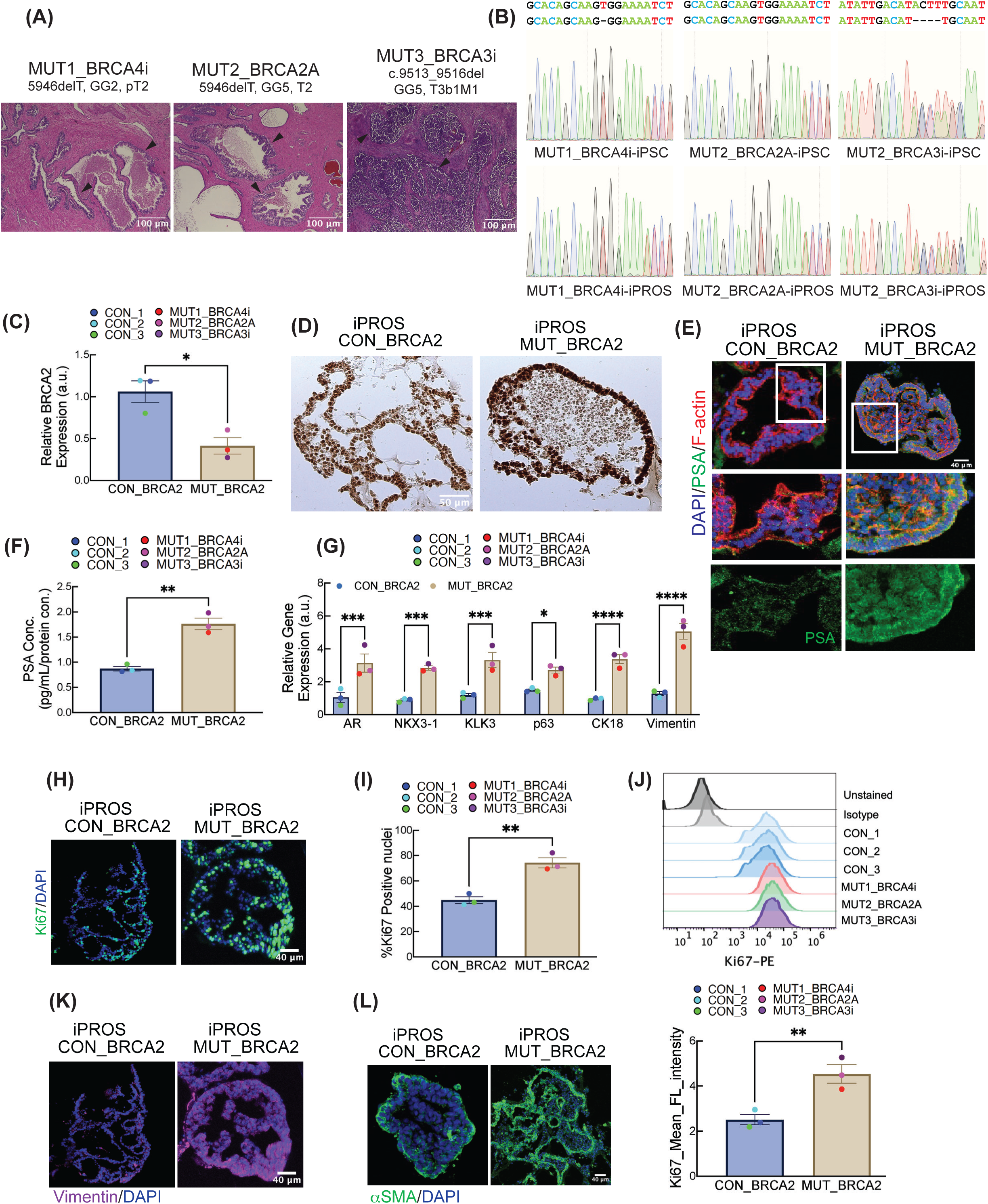
Morphological and Molecular Plasticity in iPROS with *BRCA2* mutations. (**A**) H&E staining of prostatectomy sections from 3-patients; black-arrows indicate normal/tumor sites. (**B**) Sanger-sequencing showing *BRCA2* mutations in iPSCs and iPROS. (**C**) qPCR of *BRCA2* expression in CON_ and MUT_BRCA2 iPROS. (**D**) Chromogenic images of AR. (E) IF-images of PSA and F-actin. (**F**) PSA release at week-8. (**G**) qPCR of prostate-specific genes in CON_ and MUT_BRCA2 iPROS. (**H–I**) Ki67 nuclei-index; image and quantification. (**J**) Flow cytometry of Ki67 mean fluorescence. (**K–L**) IF-images of Vimentin and α-SMA. Significance tested with pairwise comparisons (t-test with Welch correction; C,D,I,G). *p < 0.05, **p < 0.01, ***p < 0.001, ****p < 0.0001.

iPSC morphology, stemness, or proliferation was not alter by *BRCA2* mutations. No changes were observed in the fluorescence of stemness markers SOX2, OCT4, and SSE4 (**Fig. S2B**), nor in *OCT4* (**Fig. S2C**) and *SOX2* (**Fig. S2D**) gene expression. Although *BRCA2* gene expression was reduced (**Fig. S2E**), *BRCA1* expression remained unaffected (**Fig. S2F**). Ki67 nuclear staining also confirmed unchanged iPSC proliferation (**Fig. S2G,H**). These *BRCA2*-mutated iPSCs were differentiated into iPROS alongside three wild-type controls for 8 weeks. Reduced *BRCA2* expression persisted in mutant iPROS, consistent with a haploinsufficiency due to loss-of-function mutation (**Fig. 2C**).

We analyzed eight-week-old iPROS morphology using immunohistochemistry. In 72% of MUT_BRCA2 iPROS, we found AR-positive cell layers filling glandular lumens—similar to prostatic intraepithelial neoplasia, a dysplasia precursor^37^—compared to only 11% in CON_BRCA2 [pValue < 0.001] (**Fig. 2D**). MUT_BRCA2 iPROS displayed thicker, distorted luminal epithelia and increased PSA fluorescence (**Fig. 2E**), consistent with PSA gene regulation by AR and its role as a prostate tumorigenesis marker^38^. PSA release was elevated in MUT_BRCA2 (**Fig. 2F**), along with higher expression of *AR*, *NKX3.1*, *p63*, *CK18*, and *KLK3* (**Fig. 2G**).

IF analysis of MUT_BRCA2 iPROS showed increased Ki67-positive nuclei, indicating enhanced proliferation (**Fig. 2H,I**), confirmed by Ki67 flow cytometry (**Fig. 2J**). Additionally, MUT_BRCA2 iPROS exhibited cytoskeletal disorganization, loss of epithelial polarity, and diffuse localization of CK8-18 and p63 (**Fig. S2I**), with elevated and mislocalized p63 suggesting early neoplastic changes^39,40^. Signs of epithelial-mesenchymal transition (EMT) were evident through increased Vimentin (*VIM*) expression (**Fig. 2G**), and enhanced fluorescence of Vimentin and α-smooth muscle actin (SMA) (**Fig. 2K, L**), indicating an early EMT onset^41^.

### Transcriptomic profile of iPROS with germline BRCA2 mutations correlated with PCa gene expressions

Transcriptomic profiling of MUT_BRCA2 iPROS offers critical insights into molecular pathways associated with PCa development and cellular dysfunctions linked to *BRCA2* mutations^42,43^. To explore these differences, we performed total-mRNA sequencing on eight-week-old MUT_BRCA2 iPROS from three independent differentiation experiments and compared them to CON_BRCA2 total-mRNAseq datasets. Initial transcriptome analysis revealed batch effects across differentiation sets (**Fig. S3A**), commonly observed in iPSC differentiation due to stochastic variation in cell-type composition or patient-specific genetic backgrounds. After batch correction, unsupervised PCA separated MUT_BRCA2 and CON_BRCA2 iPROS along PC1 and PC4 (**Fig. 3A**), as guided by eigencore correlation plots analyzing batch, cell line, and genotype as co-variants (**Fig. 3B**). Differential expression analysis (DESeq2: fold-change >1.5, adjP < 0.05) identified 869 DEGs—330 upregulated and 177 downregulated in MUT_BRCA2 compared to CON_BRCA2 (**Fig. 3C**). Normalized expression showed altered levels signature genes implicated in early PCa; increased expression of *AR*, *FOXA1*, *EGFR, AMACR* and loss of *PTEN* and *RB1* and others^8,44,42,45^ (**Fig. 3D, S3B**). GO and KEGG geneset-enrichment-analysis on the DEGs identified activated signaling pathway_enrichment in MUT_BRCA2 iPROS, including glutathione metabolism, receptor clustering, lipid metabolism, and DNA adduct signaling, while cell adhesion, ECM anchoring, insulin resistance, and APP catabolism were suppressed—suggesting an oncogenic transcriptomic shift (**Fig. 3E,F, S3C,D**). Hallmark50_NES showed elevated Androgen Response, INF, KRAS, and MYC signaling, and loss of apical junctional signaling suggesting Pro-PCa signaling activation in MUT_BRCA2 iPROS^46^ (**Fig. 3H**).

**Figure 3:**
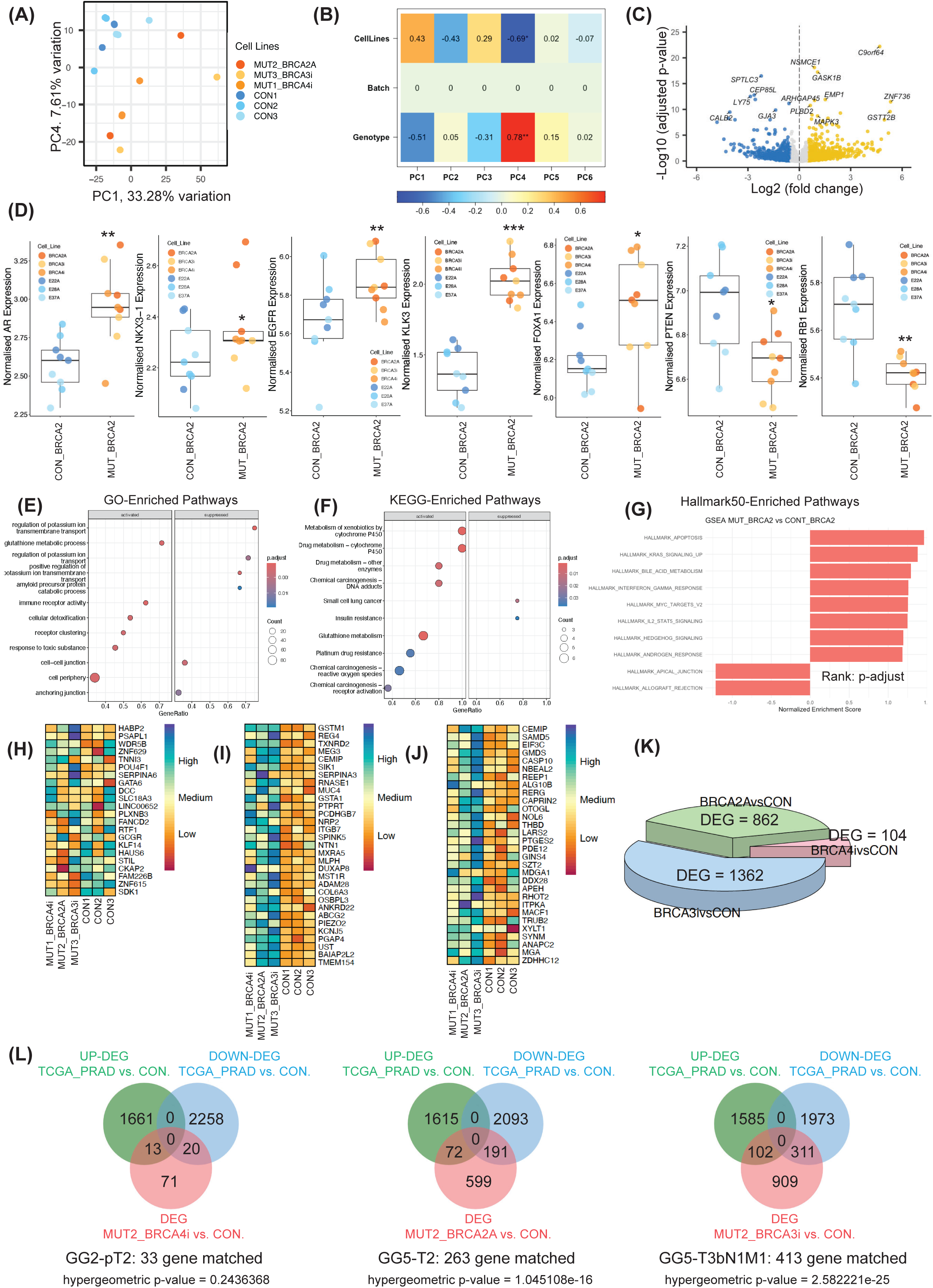
Transcriptomic characterization of iPROS with *BRCA2* mutations. (**A–B**) PCA and eigencore correlation plots showing variance among CON_ and MUT_BRCA2 iPROS. (**C**) Volcano plot of top 10 significantly altered genes. (**D**) Normalized expression of prostate- and PCa-specific genes. (E–G) GSEA analysis for GO, KEGG, and Hallmark50 pathways with significant DEGs of MUT_BRCA2 vs CON_BRCA2 iPROS. (**G**) Pie chart of transcript overlap with MSigDB PCa gene sets. (**H–J**) Heatmaps of PCa-specific genes from public-PCa-mRNAseq-datasets. (**K–L**) Pie chart showing DEGs of each MUT_BRCA2 vs CON_BRCA2, Venn diagrams comparing them with TCGA_PRAD datasets.

To assess *BRCA2* mutation-specific PCa-gene expression, we analyzed TCGA_PRAD mRNAseq-data^44^ [TCGA, Cell, 2015]. 42 genes were upregulated in *BRCA2*-mutated patients (n=5) vs Non-*BRCA2* patients (n=328), of which 22 overlapped with MUTvs.CON_BRCA2-iPROS DEGs, and 12 upregulated in MUT_BRCA2 (p-value = 1.116389e-06) (**Fig. 3H**). In TCGA-PanCancer mRNAseq-dataset^47^, 4705 DEGs were found between 494_PRAD and 51_normal patient-samples; 229 genes overlapped with MUTvs.CON_BRCA2-iPROS DEGs. 30 most upregulated genes in PRAD were similarly elevated in MUT_BRCA2 iPROS (**Fig. 3I**). Additionally, 394 DEGs were found in Primary_vs._Metastasis-PCa PDXO mRNAseq-dataset^48^, 339 of which matched MUTvs.CON_BRCA2-iPROS DEGs (p-value = 1.77289e-10), with the top 30 also upregulated in MUT_BRCA2 (**Fig. 3J**), reinforcing their oncogenic transcriptomic profile.

To evaluate patient-specific transcriptomes, we analyzed DEGs from each MUT_BRCA2 iPROS line—BRCA4i (GG2), BRCA2A (GG5), and BRCA3i (GG5 with metastasis)—against CON_BRCA2. BRCA3i, BRCA2A, and BRCA4i had 1362, 862, and 105 DEGs respectively, showing stratified gene-expression by disease stage (**Fig. 3K**). When matched these individual DEGs with TCGA_PRADvs.Normal DEGs, BRCA3ivs.CON-DEG and BRCA3ivs.CON-DEG had significant gene-overlapping with 413 and 263 respectively, whereas BRCA4ivs.CON-DEG showed only 33 gene-overlap, reflecting disease-stage-specific transcriptional fidelity (**Fig. 3L**). Finally, MUTvs.CON_BRCA2-iPROS DEG comparison was done with four PCa-genesets from MSigDB: M6698 (RAMASWAMY_METASTASIS_UP, 67-genes_upregulated in metastatic_vs_primary-PCa), M4691 (LIU_PROSTATE_CANCER_UP, 100-genes_upregulated in PCa_vs_benign-tissue), M11504 (TOMALINS_PROSTATE_CANCER_DN, 41-genes_downregulated in PCa_vs_benign-tissue), and M10319 (WALLACE_PROSTATE_CANCER_RACE_UP, 305-genes_up-regulated in PCa tissues from African-American patients compared to those from the European-American patients). For M6698, M4691, and M10319 DEGs, 36, 55, and 100 DEG-matched genes were upregulated respectively in MUT_BRCA2. Conversely, for M11504, 21 genes were downregulated in MUT_BRCA2 (**Fig. S3E**). These gene-expression similarity across these datasets (≥50% for most sets) supports a pro-PCa transcriptional environment in MUT_BRCA2 iPROS.

### Induction of tumorigenic transformation in iPROS model with dietary carcinogens

To induce PCa *in vitro* using our experimental iPROS model, we exposed 8-week-old iPROS to two dietary carcinogens—PhIP (2-Amino-1-methyl-6-phenylimidazo[4,5-b]pyridine) and MNU (N-methyl-N-nitrosourea)— for three weeks (**Fig. 4A**). Initially, were incubated them with a dose gradient (100 μM to 1 μM over 7 days) of PhIP and MNU to optimize cytotoxicity. Based on LDH cytotoxicity assays, two concentrations, 1 μM (PhIP1 and MNU1) and 5 μM (PhIP5 and MNU5), were selected for three-week exposures in iPROS with and without *BRCA2* mutations. These treatments resulted in 10–20% cytotoxicity (**Fig. S4A**). After removing the stressors at week-3, no significant differences in proliferation or PSA secretion were observed between stressed and control groups. Since carcinogenesis often shows latency, ranging from days to decades depending on carcinogen type, dosage, and genetic/immunogenic factors^49^, we extended the culture for an additional 4 weeks, assessing LDH cytotoxicity at weeks 2 and 4 (**Fig. 4B,C**). By week 4, control-iPROS treated with higher PhIP and MNU concentrations exhibited notable cell death, which was absent in *BRCA2*-mutant iPROS. Unstressed CON_ and MUT_BRCA2 iPROS maintained viability compared to stressed counterparts.

**Figure 4:**
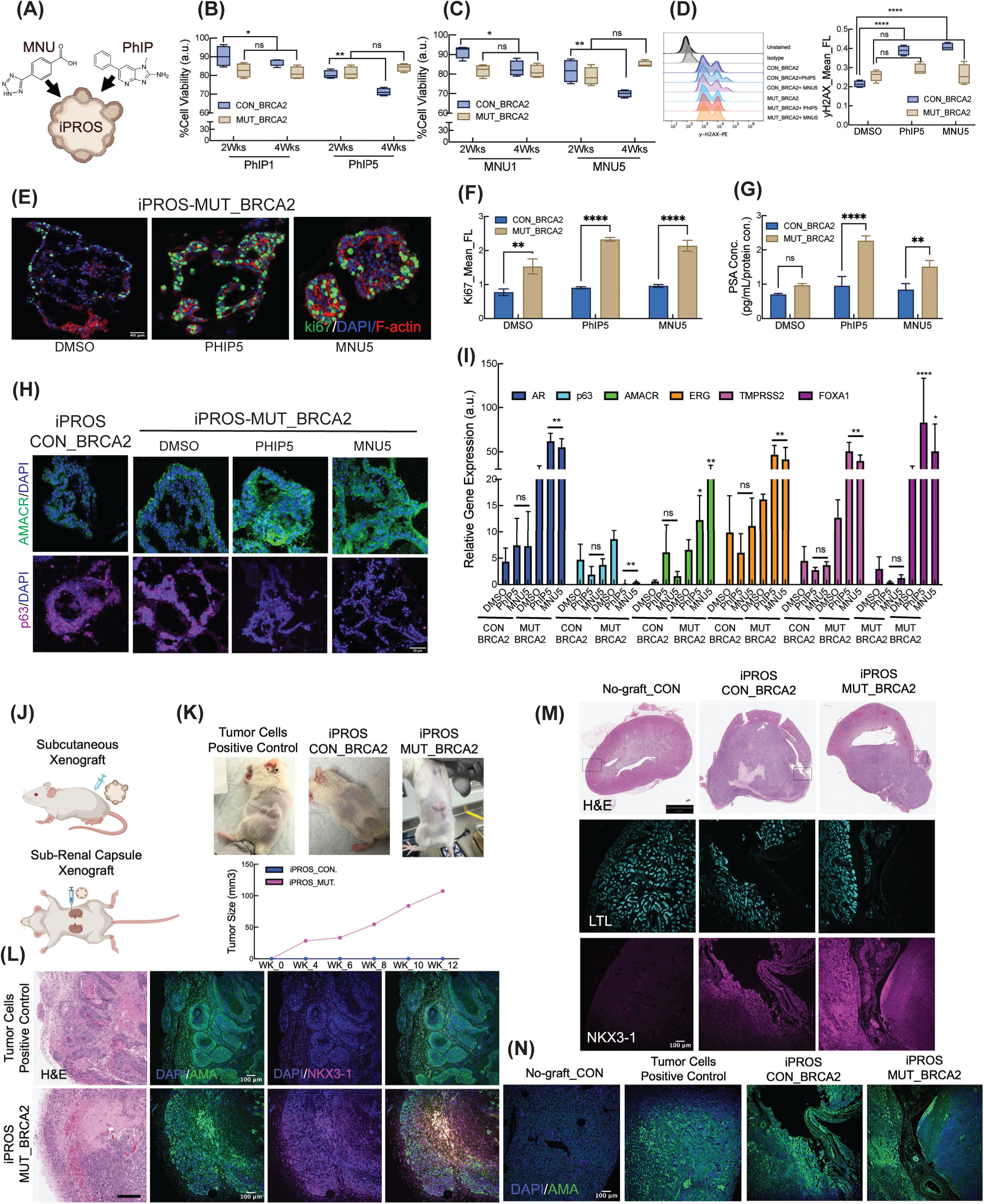
Neoplasia induction in iPROS with dietary carcinogens. (**A**) Structures of PhIP and MNU. (**B–C**) LDH assay shows cytotoxicity in CON_ and MUT_BRCA2 iPROS after 2- and 4-week of PhIP (**B**) and MNU (**C**) withdrawal. (**D**) γH2AX flowcytometry indicating DNA damage after a 3-week exposure. (**E**) Representative IF-images of Ki67-positive nuclei in 15-week-old iPROS post-treatment. (**F**) Ki67 flow cytometry after 4-week stress-withdrawal. (**G**) PSA release comparison. (**H**) Immunofluorescence for AMACR and p63. (**I**) qPCR of PCa-specific genes. (**J**) Illustration of subcutaneous and sub-renal capsules xenograft into NOD scid gamma mice (**K**) Tumor growth curve shows progressive size increase over 12 weeks post-injection. (**L**) H&E and IF-images of AMA and NKX3.1 in subcutaneous-tumors. (**M–N**) Renal capsule engraftment showing H&E and IF with LTL, NKX3.1, and AMA. Scale-bar: 100 μM (L) and significant 2-way ANOVA (multiple comparisons); *p < 0.05, **p < 0.01, ****p < 0.0001.

We hypothesized that prolonged PhIP and MNU exposure caused extensive double-strand breaks (DSBs) surpassing homologous recombination (HR) repair capacity, triggering cell death in CON_iPROS. However, MUT_BRCA2 iPROS, deficient in HR due to BRCA2 loss of function, compensate by engaging error-prone non-homologous end joining (NHEJ)^50^. Post-stress, these mutants showed reduced DSBs, evident from decreased γH2AX fluorescence (**Fig. 4D, S4B**), a sensitive DSB damage marker^51^. While NHEJ facilitates recovery, it is inaccurate, causing indels and genome instability^52^, which promotes survival and tumorigenesis^53^. This was corroborated by elevated Ki67 nuclei index (by IF) and higher Ki67 mean fluorescence (by flow cytometry) in MUT_BRCA2 iPROS treated with PhIP5 and MNU5 (**Fig. 4E,F, S4C**).

Given that PhIP and MNU induce PCa in rodents, we next examined if iPROS could mimic this tumorigenesis *in vitro*. At week 4 post-stress, PSA levels were elevated (**Fig. 4G**). Diagnostic PCa detection using tissue microarrays often employs a 3-antibody panel: AMACR (α-methylacyl coenzyme A racemase), 34βE12 (high molecular weight cytokeratin), and p63^54^. AMACR is overexpressed in PCa^55^, while loss of 34βE12 and p63 supports diagnosis. Mutant iPROS treated with PhIP5 and MNU5 exhibited increased AMACR staining and reduced p63, suggesting PCa-like features (**Fig. 4H**). RT-qPCR showed upregulation of *AR*, *AMACR*, *ERG*, *TMPRSS2*, and *FOXA1*, and downregulation of *p63*^56,57,58^ (**Fig. 4I**), indicating tumorigenic transformation in MUT_BRCA2 iPROS.

Upregulated AR and its downstream genes suggest AR-driven oncogenesis. Accordingly, PhIP5- and MNU5-stressed organoids were treated with 5 μM enzalutamide, an AR inhibitor^59^, for 2 weeks post-stress. This treatment elevated cytotoxicity and reduced proliferation in MUT_BRCA2 iPROS (**Fig. S4D,E**). Additionally, 10 μM Olaparib, a PARP inhibitor inducing synthetic lethality by targeting NHEJ in MUT_BRCA2 iPROS cells^60^, caused selective cytotoxicity and increased γH2AX staining these iPROS (**Fig. S4F,G**). These findings affirm the iPROS model as a reliable human-relevant *in vitro* platform for studying prostate tumorigenesis.

### Xenotransplanted BRCA2 mutant iPROS in immunodeficient mice promote tumorigenesis

Hayflick limits the long-term expansion of organoids *in vitro*, making it difficult to fully model tumorigenesis^32^. Moreover, tumor-stromal interactions and vascularization are critical for tumor progression, maturation, and metastasis^61^. iPROS cultures ceased expanding after 14–16 weeks and began to die (data not shown); with PhIP and MNU 3 weeks treatment, this extended up to 18-22 weeks before they die out. To support further growth in a host environment with a more favorable tissue microenvironment, we conducted a xenograft study by injecting iPROS either subcutaneously or into the sub-renal capsules of 6–8-week-old NOD scid gamma mice (**Fig. 4J**). We injected CON_ and MUT_BRCA2 iPROS, a human tumor cell line (positive control, human-urinary-bladder cancer cell-line 5637), and Matrigel alone. The renal capsule is an advantageous ectopic site for prostate tissue xenografting^62,15^.

Four weeks post-injection, large tumors formed in mice receiving the tumor cell line subcutaneously, and three small tumors developed in MUT_BRCA2-iPROS-injected mice (**Fig. 4K**). Subcutaneous tumor growth curves were tracked for 12 weeks, after which mice were euthanized and tumors collected. The positive control mouse reached a tumor-size of 300 mm³ by week 6 and was euthanized earlier. No tumors formed in CON_BRCA2-iPROS injected mice. Alongside hematoxylin & eosin (H&E) staining (**Fig. 4L, S4H**) we identified the integration of both human-tumor cells and iPROS cells into the mouse subcutaneous tumor using human-specific anti-mitochondria antibody (AMA, 113-1), which stains explicitly human mitochondria (a specifc “spaghetti-like” staining) and does not cross-react with mouse or rat tissues (**Fig. 4L**)^63^. Only MUT_BRCA2 iPROS tumors expressed NKX3.1, a prostate-specific marker. In sub-renal capsule xenografts, LTL-negative^64^ (kidney marker), but NKX3.1 positive, large-mass outgrew only in MUT_BRCA2 iPROS and positive control-tumor cell injected kidneys (**Fig. 4M, S4H**). PCa markers ERG and PSMA fluorescence were elevated in MUT_BRCA2 iPROS-transplanted kidneys (**Fig. S4M**), supporting *in vivo* tumorigenesis by MUT_BRCA2 iPROS. Positive control, CON_ and MUT_BRCA2 iPROS transplated kidney sections were positive for AMA confirming human prostate cell integration (**Fig. 4N**).

## DISCUSSION

Patients’ iPSC-derived organoids are emerging as valuable tools for modeling organ development and disease progression, including cancers^65^. While prior models with human iPSCs showed prostate-like organoids, these were reliant on co-culture with rodent UGM^14,15^, reducing their utility in pre-clinical studies. In this study, we developed a pre-clinical, rodent cell–free prostate organoid (iPROS) model that reliably recapitulates human prostate organ morphology, gene expression, and function^12^. iPROS with *BRCA2* risk variants displayed functional implications in morphological and molecular plasticity during prostate tumorigenesis^66^. Our results demonstrate the potential of iPROS as an effective platform for *BRCA2*-related PCa risk prediction, personalized biomarker evaluation, and therapeutic screening. iPROS differentiation from iPSCs followed a well-characterized protocol mimicking prostate glandular development^26^, enabling detailed mapping of epithelial-mesenchymal interactions from definitive endoderm through hindgut, prostate progenitor, and epithelial budding stages—circumventing the limitations of adult tissue-derived organoids.

The use of defined small molecules and pathway modulators enabled generation of 3D prostate organoids mimicking prostate lobes and ducts, showing both basal and luminal epithelial structures. iPROS expressed key prostate markers including Androgen Receptor (AR), NKX3.1, and p63, critical for prostate development. PSA secretion increased over time indicating androgen responsiveness and functional maturation. Detection of PSA confirmed the ability of iPROS to recapitulate key physiological features *in vitro*. Importantly, iPROS reproduced human prostate-like architecture, including luminal, basal, and neuroendocrine epithelial lineages, as confirmed by IF/Immunohistochemistry showing CK8-18, p63, and Chromogranin A expression. Transcriptomic analysis further validated physiological relevance by aligning closely with human adult prostate tissue gene profiles.

Pathogenic *BRCA2* mutations are common in aggressive, metastatic PCa with high-GG tumors. We derived iPSCs from three patients carrying pathogenic *BRCA2* mutations with different GG and stages, and differentiated them into iPROS. This enabled us to study the impact of germline *BRCA2* mutations on organoid development and tumorigenesis^10^. As a tumor suppressor, *BRCA2* maintains DNA integrity during cell division. Its mutation impairs DNA repair, leading to genomic instability and uncontrolled proliferation. These deficiencies can disrupt other genes linked to growth and survival, promoting tumorigenesis^67^. Our findings support this: *BRCA2* mutations aligned with altered prostate epithelial morphology and gene expression but did not affect iPSC proliferation or stemness, suggesting their influence manifests post-differentiation.

Increased Vimentin and SMA levels, along with disrupted epithelial polarity in MUT_BRCA2 iPROS, are consistent with EMT transition, indicating predisposition to tumorigenic transformation. Transcriptomic profiling revealed enrichment of PCa signature genes in MUT_BRCA2 iPROS. GSEA showed pro-oncogenic AR, MYC, MAPK, and Hippo signaling upregulation, cell-peripheral signaling, cell-cell adhesion loss, and extracellular matrix (ECM) remodeling, all hallmarks of prostate-epithelial tumorigenesis^42,68^. MUT_BRCA2 iPROS shared significant transcriptomic overlap with prostate adenocarcinoma (PRAD), further supporting its relevance as a disease model. The varying degrees of transcriptomic similarity between different GG-MUT_BRCA2 iPROS and TCGA_PRAD datasets underscore the iPROS model’s potential to study primary and advanced PCa in a patient-specific context^68^. Finally, *in vivo* xenotransplantation confirmed higher tumorigenic potential of *BRCA2*-mutated iPROS in both subcutaneous and renal capsule microenvironments.

We found that PhIP and MNU, cause genotoxic stress and induce PCa in rodent models^20,21^, also induced DNA damage in iPROS. Despite cytotoxic stress, unrepaired or mismatch-repaired DNA allowed MUT_BRCA2 iPROS to survive, intensifying their oncogenic potential. Importantly, the carcinogen-treated iPROS responded like human PCa: elevated expression of PCa-specific genes (*AR*, *AMACR*, *ERG*, *TMPRSS2*) and loss of the basal marker p63 indicated onset of aggressive-PCa transformation. Furthermore, iPROS with *BRCA2* mutations exhibited increased sensitivity to AR inhibitor enzalutamide and PARP inhibitor olaparib, confirming their transformation state and dependence on AR signaling and DNA repair pathways.

In conclusion, we developed patient-specific prostate organoids (iPROS) that mimic human prostate morphology and function. iPROS with pathogenic-*BRCA2* mutation forms tumors *in vivo*, and show dietary-carcinogen vulnerability, providing a robust preclinical platform to study mutation-specific-PCa and advance precision oncology therapies.

## ACKNOWLEDGEMENTS

The authors thank Dr. Wong-Valencia for help with iPROS dissociation, Fangyuan Qu and Yongqi Lin for mouse care. Dr. Sunyoung You for discussing RNAseq data analysis, Dr. Soshana Svendsen for critical reading of the manuscript. This work was supported by an award from the Urological Research Foundation to CNS and DT, and institutional support to BRA (Donna and Jesse Garber Award for Cancer Research, 2025) and CNS from Cedars Sinai Mecical Centre.

## AUTHOR CONTRIBUTIONS

CNS. And DT. conceived project. BRA conceptualized/developed iPROS and oncogenesis model, designed experiments, and wrote manuscript with CNS. BRA, CNS, DT, and WJC edited manuscript. BRA performed experiments with GL, PA, PA, MGO, YF, SG, and ZM. BRA, MJW, and SB analyzed RNAseq data.

## DECLARATION OF COMPETING INTERESTS

Authors declare a patent application filling related to this work.

## SUPPLEMENTAL INFORMATION

Document S1: Key resources table, Supplementary-Figure Legends, Materials and Methods

Table S1: iPSCs details.

## STAR METHODS

### KEY RESOURCES TABLE

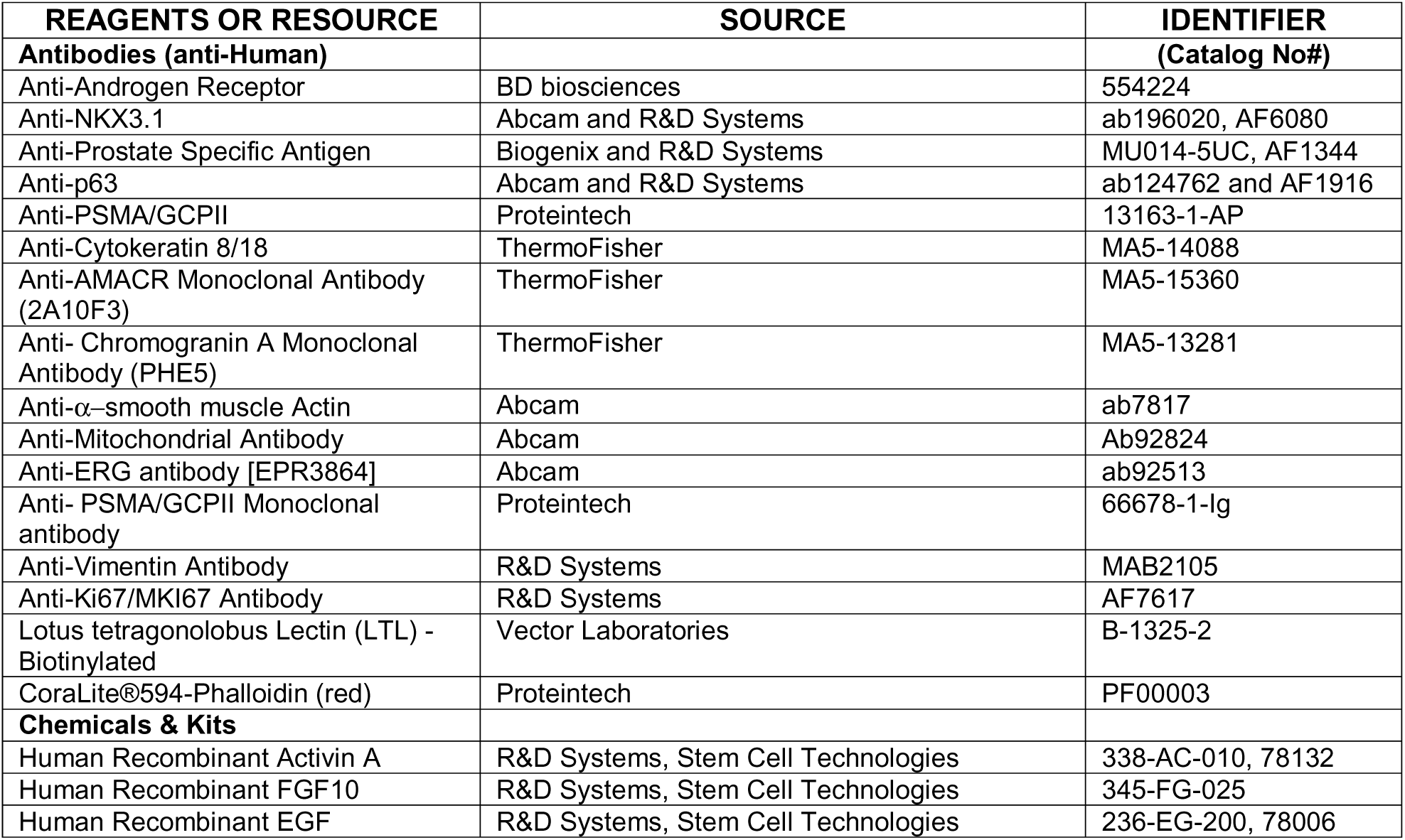
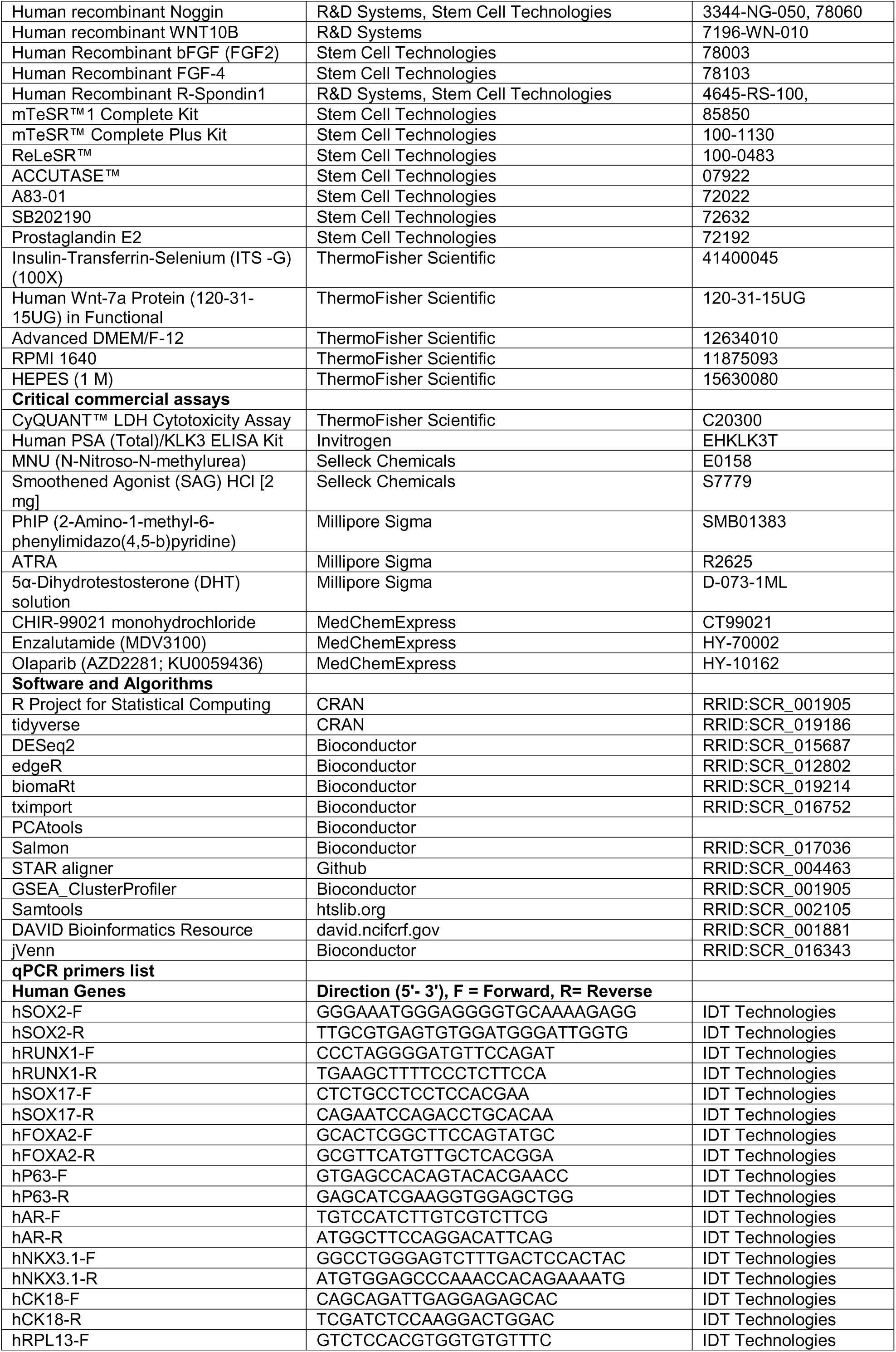
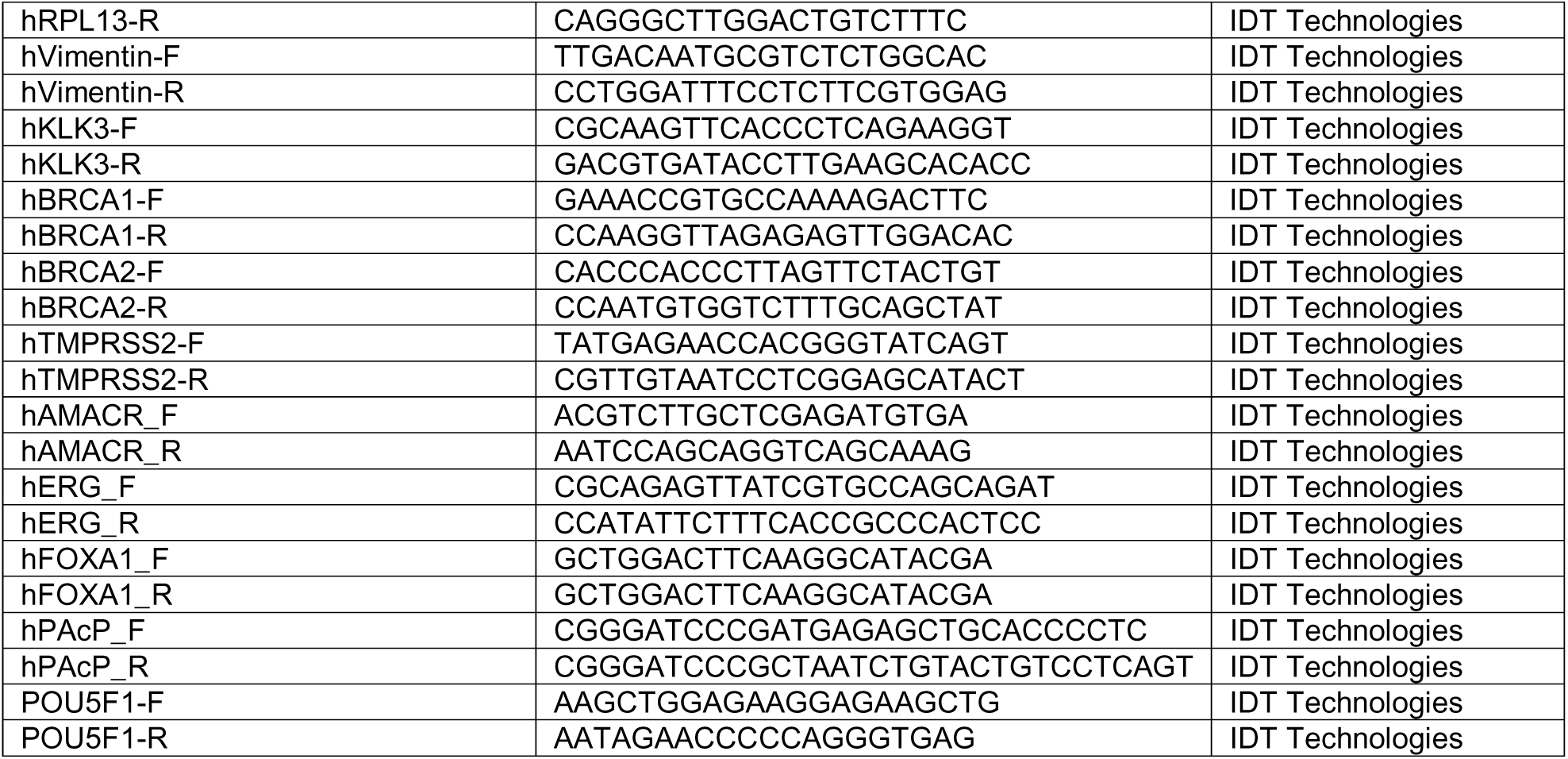

### RESOURCE AVAILABILITY

#### Lead contact

Further information and resource requests should be directed to the lead contact, Clive Svendsen.

#### Materials availability

The iPSC lines used in this study can be searched and selected through the catalog at the Cedars-Sinai Biomanufacturing Center (https://biomanufacturing.cedars-sinai.org) for order fulfillment.

#### Data and code availability

All datasets generated during and/or analyzed during the current study and the R-codes are available upon request.

### SUPPLEMENTARY TABLE

**Table S1:** List of lines with identifier and other details used in the current study.

| iPSC Line | Mutations | Code used in Manuscript | Age at PBMC Draw | Prostate Cancer Diagnoses | Gleason Grade Group (ISUP) | Path Stage if available | Lymph Node Metastases | Distant Metastases | Curr. Status | Cancer aggressiveness |
| --- | --- | --- | --- | --- | --- | --- | --- | --- | --- | --- |
| 0002iBRCA2 | c.5946delIT | BRAC2A | 68 | 1 | 5 | T2 | 0 | 0 | NED | High |
| N0U3iPRC | c.9513_9516del | BRAC3i | 59 | 1 | 5 | T3bN1M1 | 1 | 1 |  | High |
| C0S4iPRC | c.5946delIT | BRAC4i | 70 | 1 | 2 | pT2 | 0 | 0 | NED | Low |
| CSEDI022iCRT | NA | CONT.1 | 79 | 0 | NA | NA | NA | NA | NA | NA |
| CSEdi028CRT | NA | CONT.2 | 79 | 0 | NA | NA | NA | NA | NA | NA |
| CSEdi037iCRT | NA | CONT.3 | 80 | 0 | NA | NA | NA | NA | NA | NA |

### MATERIALS AND METHODS

#### iPS Cell Lines

All iPSC lines with *BRCA2* mutations were generated at the iPSC Core at Cedars-Sinai Medical Center. Patients peripheral blood mononuclear cells (PBMCs) were transfected with a non-integrating episomal plasmid expressing seven factors: OCT4, SOX2, KLF4, L-MYC, LIN28, SV40LT, and p53 shRNA (pEP4 E02S ET2K, pCXLEhOCT3/4-shp53-F, pCXLE-hUL, and pCXLE-hSK). All cell lines and protocols in this study were conducted in compliance with the guidelines approved by the Stem Cell Research Oversight Committee (SCRO) and Institutional Review Board (IRB) under IRBSCRO Protocols Pro00032834 (iPSC Core Repository and Stem Cell Program) and Pro00021505 (Svendsen Stem Cell Program). Three existing male control iPSC lines with wild-type *BRCA2* were selected from the Cedars-Sinai Biomanufacturing Center iPSC Core. These control lines were reprogrammed from healthy donor PBMCs, namely CSEDi022A, CSEDi028A, and CSEDi037A. The presence of *BRCA2* heterozygous mutations in these iPSC lines was confirmed through DNA sequencing analysis. *BRCA2* mutations identified were specific to each patient line: CSN0U3iPRC (c.9513_9516del, located in exon 11), CS0002iBRCA2 (c.5946delT, located in exon 19), and CSC0S4iPRC (c.5946delT, located in exon 19). These mutations aligned with the patients’ clinical diagnoses.

#### iPSC Culture

Control and *BRCA2*-mutated iPSCs were cultured in mTeSR®Plus medium on growth factor-reduced Matrigel™ Matrix (BD Biosciences)-coated plates at 37°C in a 5% CO2 incubator. When human iPSC colonies reached 70–90% confluence, they were washed with Versine and gently lifted with ReLeSR (STEMCELL), and then replated at a 1:6 ratio. All cell lines were tested for mycoplasma contamination monthly.

#### Differentiation of iPROS from iPSCs

iPROS differentiation from iPSCs involved three major steps: iPSC to Definitive Endoderm (DE) differentiation, followed by differentiation hindgut endoderm (HG) and prostate progenitor specification in 3D culture, and finally the induction of AR signaling in progenitor population for further differentiation.

##### DE Differentiation (3-day Protocol)

iPSCs growing in mTeSR®Plus medium until DE induction. In a confluent well of a 6-well plate (approximately 1.2×10^6 cells), cells are dissociated using Accutase ^63^ in mTSer1 media containing 10 μM Y-drug and plated in a 24-well plate pre-coated with Matrigel (0.25-0.5 mg/ml). On Day 1, the medium is replaced with 1 mL of DE media (RPMI, 1x P/S, 1x L-glutamine, 3 μM Chir99203, and 100 ng/ml Activin). On Day 2, the media is changed to a fresh Day 2 formulation (RPMI, 1x Pen/Strep, 0.2% FBS, 100 ng/ml Activin A), and on Day 3, the media is switched to Day 3 media (RPMI, 1x Pen/Strep, 2% FBS, 100 ng/ml Activin).

##### Hindgut Endoderm & Prostate Progenitor Differentiation (2+2 Days Protocol)

Following DE differentiation, the media is replaced with HG media for 4 days, refreshing the media daily. On Days 1 and 2, cells are exposed to AdvDMEM/F12 (with 200 mM L-glu, 1x P/S, 15 mM Hepes) supplemented with 2% FBS, 500 ng/ml FGF4, and 500ng/ml WNT3B. On Days 3 and 4, the media is changed to a formulation containing 2% FBS, 200 ng/ml FGF4, 500 ng/ml FGF10, and 500 ng/ml WNT10B. At this stage, 3D aggregates form beneath the monolayer, which are collected by scraping, spun at 200g for 3 minutes, and preferably separated as 3D aggregates for subsequent steps.

##### Inducing of AR Signaling

Matrigel is prepared by adding 20 μl of 1x B27, 1.5 μl of 100 μg/ml EGF, and 1.5 μl of 100 μg/ml Noggin to 1 ml of stock Matrigel (∼10 mg/ml). The cell pellet is mixed 1:1 with this Matrigel cocktail, with each drop of the mixture not exceeding 70 μl (ideally 50 μl) per well of a 24-well plate. After 3-4 minutes in the hood, the plate is flipped upside down, allowing the cells to hang in the Matrigel drop for 9 minutes in the incubator. Following this, 500 μl of media is added to each Matrigel dome well. For the next 5-7 days these prostate progenitors were cultured in media consisting of AdvMEM12 (with 200 mM L-glu, 1x P/S, 15 mM Hepes), 1x B27, and 2% ITS, along with 500 ng/ml R-Spondin1, 100 ng/ml Noggin, 100 ng/ml EGF, 10 nM ATAR, 1.5 μM DHT, 2 μM CHIR-99021, 10 nM SAG (SHH agonist), and 100 ng/ml FGF10. Media should be replaced every third day.

##### Long-term iPROS Culture

From Week 2 onwards, organoids are maintained in a modified media formulation: AdvMEM12 with 200 mM L-glu, 1x P/S, 15 mM Hepes, 1x B27, 2% ITS, 1.25 mM N-acetylcysteine, 1 μM prostaglandin E2, 10 mM nicotinamide, 0.5 μM A83-01 (TGFβ/Smad inhibitor), 10 μM SB202190 (p38 MAPK inhibitor), 500 ng/ml R-Spondin1, 100 ng/ml Noggin, 100 ng/ml EGF, 100 ng/ml FGF2, 100 ng/ml FGF10, 10 nM ATAR, 10 nM DHT, 10 nM SAG, 2 μM CHIR-99021, and 10 μM Y-drug. After Week 3, Y-drug and ATAR are removed, but they should be reintroduced when organoids are split. Organoids were passaged every 2-3 weeks. To passage, organoids are dissociated with Express-TrypLE to single cells, counted, and mixed with a 1:1 Matrigel cocktail to regrow organoids with even cell numbers. This method ensures a consistent number of organoids per dome.

##### Cryopreservation and Revival

For cryopreservation, Matrigel is removed, and Cryostor media is added to the organoids. After mechanical disruption with a pipette, the organoids are frozen with Cryostor cell freezing media (STEMCELL). For the revival, Cryostor is replaced with the 1:1 Matrigel cocktail, and after one week, organoids can be re-split and counted to ensure even distribution.

#### RNA Extraction and Quantitative PCR Analysis

Total RNA was extracted from cells using the QIAGEN Rneasy Mini Kit, following the manufacturer’s instructions. One microgram of the purified RNA was then used to synthesize cDNA using the Quantitect Reverse Transcription Kit (QIAGEN). Quantitative real-time PCR was carried out with SYBR Select Master Mix (Applied Biosystems). Gene expression levels were normalized to *GAPDH* and *RPL13* housekeeping genes expressed as fold changes relative to control samples. All experiments were conducted in triplicate, and the primers used are listed in the reagent table. At least three independent experiments were performed for all genes with three technical repeats.

#### Fluorescence-Immunohistochemistry of iPROS and Immunocytochemistry of iPSC Cells

iPROS were fixed in 4% paraformaldehyde (PFA) in phosphate buffered saline (PBS; with Ca2+ and Mg2+) for one hour at room temperature (RT), then rinsed three times with PBS. After fixation, they were cryoprotected overnight in 30% sucrose at 4°C and embedded in OCT compound (Tissue-Tek). Frozen sections, 10 µm thick, were cut using a cryostat, mounted on glass slides, and stored at –20°C. Before staining, sections were rehydrated, permeabilized, and blocked with PBS with 0.5% Triton X-100 and 10% normal human serum in PBS for one hour at room temperature. Sections were then incubated overnight at 40C with primary antibodies diluted the blocking solution containing 0.05% Triton X-100 (PBS-T). The secondary antibody was diluted (1:1000) with 5% Normal Donkey Serum in PBS-T for 1 hour. iPSCs were fixed in 4% PFA in PBS (with Ca2+ and Mg2+) for 20 min at RT. This was followed by permeabilization for 5 minutes with 0.25% Triton X-100, then blocking with 5% Normal Donkey Serum in PBS-T for two hours in RT, and then the secondary antibody was diluted (1:1000) in 5% Normal Donkey Serum in PBS-T for 1 hour. Nuclei were counterstained with DAPI (4’,6-diamidino-2-phenylindole). Finally, the slides were mounted with antifade mounting media and dried before imaging with a Nikon-Ti Confocal microscope. Each image represents at least three independent experiments. The primary antibodies used in this study are listed in reagent table.

#### iPROS Xenografts

All procedures involving animals and their care were approved by the Institutional Animal Care and Use Committee of CSHS (IACUC008253) in accordance with institutional and National Institutes of Health guidelines. Male immunodeficient (NGS) mice (n=1-2/group), 6-8 weeks old were injected, either subcutaneously into the flanks or underneath the renal capsule, bilaterally as follows. Each administration site received 1 million (1 × 10^6 cells) positive control tumor cells (positive control group), matrigel alone (negative control group) or iPSC derived organoids [200 organoids (∼150-300 mm in diameter, consists of ∼ 2 million single cells, uninformedly sheared with 1 ml pipette tips,) mixed with cold Matrigel (50:50)] for CON_BRCA2 and MUT_BRCA2 groups. Briefly, animals were anesthetized with induced with isoflurane (1-3%) and maintained on a nose cone, placed on a heating pad (37°C) ear tagged, skin was shaved and aseptically prepped using betadine and 70% alcohol. For subcutaneous delivery, a single puncture hole was used to deliver the cells into the subcutaneous tissue of both flanks via a 18-gauge needle. For subrenal capsule delivery, a ∼1cm midline incision was made in the back in between the kidneys, the incision was moved over the flank, a small incision was made in the muscle, and the kidney gently exposed through the incision. The kidney was kept hydrated using 0.9% sterile saline. A small incision was made in the kidney capsule using a 23-gauge needle, using an elevator the capsule was separated from the kidney to create a small pocket. The cells or iPROS were delivered in 10-50µL of Matrigel using PE-50 tubing. The kidney was gentrly placed back into position through the incision, the muscle layer sutured with absorbable 5-0 suture and the skin incision closed with wound-clips. Buprenex (0.1mg/kg) and Carprofen (5mg/kg) were given for analgesia post-operatively. Animals were monitoried regularly for signs of tumor growth. Mice were euthanized 6 months after the graft implant or when the tumour size reached ≥300-500 mm^3^.

#### Immunohistochemistry of Xenograft iPROS Tissues

Dissected xenografted tissue was fixed in 10% formalin for 1 hour at room temperature, then overnight at 4°C, washed with PBS, and stored in 70% ethanol overnight. Samples were processed by the Cedars-Sinai Biobank Core for paraffin embedding and sectioned at 5 µm. Slides were deparaffinized with xylene (3x, 10 min each), then rehydrated through graded ethanol solutions (100%, 95%, 75%, 50%) for 5 minutes each. After rinsing twice with tap water, antigen retrieval was performed by microwaving in Vector Unmasking Solution (citric acid-based) and then cooling for 30 minutes.

Endogenous peroxidase activity was blocked with 0.3% H_2_O_2_ in methanol for 30 minutes. Slides were blocked with 3% BSA in PBS-T for 1 hour, then incubated overnight at 4°C with primary antibody (1:200). After PBS washes, sections were treated with a biotinylated secondary antibody, followed by ABC reagent and DAB development. Slides were counterstained with hematoxylin, dehydrated, cleared in xylene, and mounted with coverslips. All H&E staining of original patient tissues and xenografted iPROS tissues was done in the biobank core. The slides were stained with hematoxylin (ImmunoMaster Hematoxylin, American MasterTech Scientific, Inc.) for 8 minutes, followed by a 5-minute rinse under running tap water. They were then briefly stained with eosin (Eosin Y Phloxine B, American MasterTech Scientific, Inc.) for 10 seconds and rinsed again with tap water. Afterward, the slides were dehydrated through a graded ethanol series—50%, 75%, 95%, and 100%— and cleared in xylene, each step lasting 1 minute. Finally, the tissues were mounted with a glass coverslip using Richard-Allan Scientific Mounting Medium.

#### Ki67 Positive Nuclei Count by Image Segmentation and Particle Analysis

iPROS sections or iPSCs were stained for Ki67 using a standard immunofluorescence protocol, and nuclei were counterstained with DAPI. Fluorescence images were acquired at 20x magnifications to ensure optimal resolution of Ki67-positive nuclei. Images were processed using Fiji-ImageJ. First, they were converted to 8-bit grayscale, background noise was reduced by applying a median filter (radius 2-3 pixels). After binary water shading, the fluorescence signal corresponding to both DAPI and Ki67 staining was thresholded using the “Threshold” tool to musk the nuclei and exclude non-specific background staining, ensuring accurate detection of the Ki67-positive signal. Then, using the “Analyze Particles” function in Fiji-ImageJ, the segmented DAPI and Ki67-positive areas were further analyzed. The parameters for particle analysis were set to 100 to 1500 mm2, and circularity 0.4-1 which exclude artifacts and only count particles of a defined size range corresponding to individual nuclei. The “Show Results” option provided a count of both DAPI and Ki67-positive nuclei in the image. The number of Ki67-positive cells was expressed as a percentage of total DAPI-positive nuclei to account for variations in cell density. Statistical Analysis: Data from multiple images (n ≥ 3 per condition and at least six segments) were used for statistical analysis. Ki67-positive cell counts were normalized to the total number of DAPI-positive nuclei per image, and the results were presented as the mean percentage of Ki67-positive nuclei ± SEM.

#### mRNA-seq Experiment and Analysis for Transcriptional Profiling of iPROS

Total RNA was extracted from samples using the QIAGEN Rneasy Mini Kit according to the manufacturer’s instructions. RNA concentration and purity were measured using a NanoDrop spectrophotometer, and RNA integrity was assessed using the Agilent 2100 Bioanalyzer. Only RNA samples with an RNA Integrity Number (RIN) greater than 8 were used for sequencing. Standarised cDNA library preparation (poly A enrichment) was done as suggested by manufacturer, using the Illumina TruSeq Stranded mRNA Library Preparation Kit. Libraries were quantified using a Qubit fluorometer and assessed for quality on the Bioanalyzer. Pooled libraries were sequenced on an Illumina NovaSeq X Plus (PE150) platform with pair-end reads at a depth of ∼25 million reads per sample. Base calling and demultiplexing were performed using Illumina bcl2fastq software. Raw reads were quality-checked using FastQC and trimmed using Trimmomatic to remove adapters and low-quality bases.

High-quality reads were pseudoaligned to the GRCh38.p13 human reference genome using Salmon (v1.4.0) with default parameters. Transcript-level abundances were imported and summarized to the gene level using the R package *tximport*. Genes with an average read count below 3 across all samples or lacking HGNC annotation were excluded. Normalization and differential expression analysis were conducted using the DESeq2 package in R. Raw counts were transformed using the variance stabilizing transformation (VST) for downstream analysis and visualization. Gene expression patterns, heatmaps were generated using the *tidyverse* and *ggplot2* R package. Principal component analysis (PCA) was used to assess sample clustering and variance across conditions. Gene symbols were added via the addIDs() function. All rows without a corresponding gene symbol were removed. All genes with total counts less than the number of samples were removed from subsequent analysis. Samples were processed and analyzed using DESeq2. Gene expression was normalized using variance stabilizing transformation, and batches were corrected using limma::removeBatchEffect. Volcano plots were generated using the Enhanced Volcano package and PCA was performed with PCAtools in R. Gene set enrichment analysis was conducted with Clusterprofiler with GO and KEGG biological process gene set. Differential gene expression analysis was performed using DESeq2 in R. Counts were normalized using DESeq2’s median-of-ratios method, and differentially expressed genes were identified based on an adjusted p-value < 0.05. Data visualization, including principal component analysis (PCA) and heatmaps, was conducted using the *ggplot2* package. All plots were generated in R (version 4.2.0). Gene Set Enrichment Analysis (GSEA) was performed using the pre-ranked method in the GSEA_ClusterProfiler2.R and GSEA_preranked_list.R, querying the MsigDB collections. Enriched pathways and gene sets were considered significant at a false discovery rate (FDR) of < 0.05.The whole pipeline, including code and parameters, is available upon request. For, Venn diagram building, Jvenn software were used^69^.

#### PSA ELISA

iPROS culture medium was collected at indicated time points and centrifuge at 1,000 x g for 10 minutes to remove any cellular debris. Store the supernatant at −80°C until further analysis. We used Human PSA (Total)/KLK3 ELISA kit from Thermo-Fisher, and prepared reagents according to the manufacturer’s instructions, including PSA standards, wash buffer and detection reagent. Add 50 µL of each standard and 50 µL of the sample supernatant to the corresponding wells in the ELISA plate. Include blanks and controls as needed. The plate was incubated at room temperature (RT) for 2-3 hours to allow antigen-antibody binding. Wash the plate three times with wash buffer to remove unbound substances. Add 100 µL of enzyme-conjugated detection antibody added to each well and incubate for 1 hour at RT. After washing, substrate solution was added and incubated for 15 minutes. Finally, we added the stop solution as recommended. Absorbance was measured at 450 nm using a microplate reader, constructed a standard curve using the known PSA standards, and calculated the concentration of PSA in the samples. Samples were normalized based on total protein concentration using a BCA assay and represented as PSA release per µg of protein.

#### Lactate Dehydrogenase (LDH) Cytotoxicity Assay

Cell membrane integrity and cytotoxicity were assessed using a LDH release assay, which was performed using the CyQUANT-LDH Cytotoxicity Assay Kit (ThermoFisher, USA.) according to the manufacturer’s instructions. Briefly, ∼200 iPROS were embedded in matrigel and allowed to adhere for 2 days before compound treatment. Following treatment with test compounds at indicated concentrations and time points, 25 µL of the culture supernatant from each well was transferred to a new flat-bottom 96-well plate. An equal volume (25 µL) of LDH reaction mixture was added to each well and incubated at room temperature in the dark for 30 minutes. The enzymatic reaction converts lactate to pyruvate, resulting in the reduction of a tetrazolium salt to a red formazan product. The absorbance was measured at 490 nm, with a reference wavelength of 680 nm, using a microplate reader. Cytotoxicity (%) was calculated using the following formula:

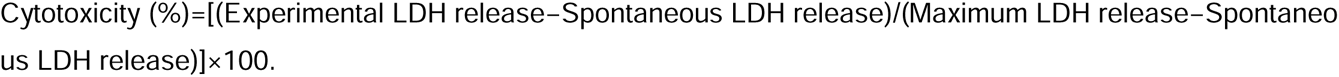

Spontaneous LDH release is when cells/organoids are incubated with medium only, and Maximum LDH release is obtained when iPROS were treated with lysis buffer provided in the kit. All experiments were performed in triplicate and repeated at least three times independently. Data are expressed as mean ± SEM.

#### Statistical Analysis

Statistical analyses were conducted using Prism software (GraphPad Software, La Jolla, California). Quantitative data are presented as mean values ± Standard Error of the Mean (SEM) and analyzed using student t-test with Welch correction or with one-way or 2-way ANOVA (for unequal variances) with multiple pair comparisons; analysis was done across three biological replicates, if not it is otherwise mentioned in figure legends. Statistical significance was determined with *p ≤ 0.05, **p ≤ 0.01, ***p ≤ 0.001, and ****p ≤ 0.0001.

### SUPPLEMENTRAY FIGURE LEGENDS

**Figure S1:**
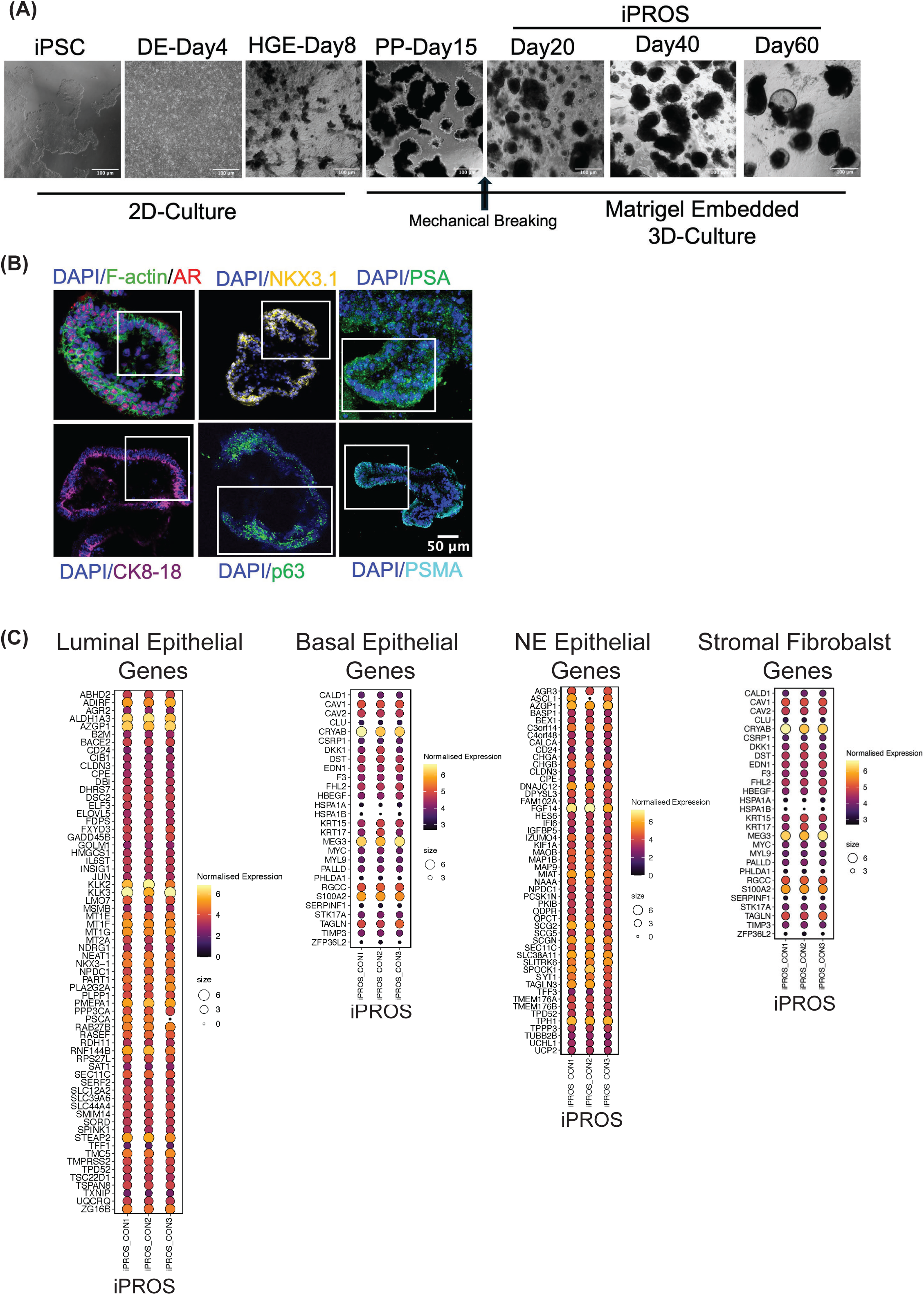
(**A**) Evos-Brightfield images showing the stages of iPROS differentiation. (**B**) Represented immunofluorescence images showing prostate tissue-specific makers in iPROS: AR, NKX3.1, CK8/18, p63, PSA, and PSMA with DAPI counterstain, insets are separately shown in Fig.1D. (**C**) Heatmaps showing the TPM normalized expression of genes in iPROS. Genelist generated from Human prostate epithelial (Luminal, Basal, and NE) and stromal cell-specific transcripts. The top 100 highly expressed genes were selected from each cell type for comparison.

**Figure S2:**
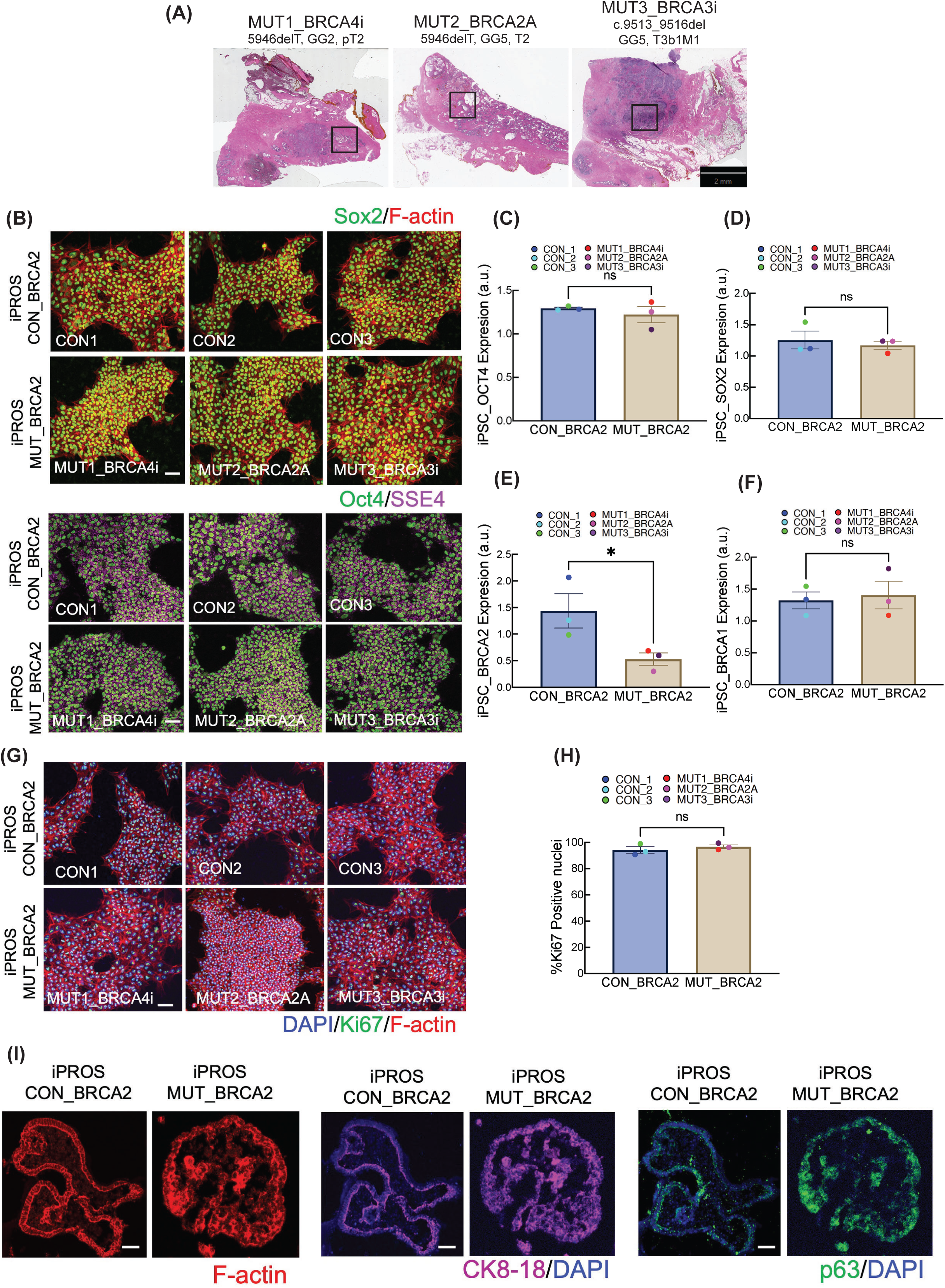
(**A**) H&E staining of prostatectomy sections from 3 patients; full slide image, insets are separately shown in Fig.2A. (**B**) Representative images showing the Sox2, Oct4, and SSE4 immunostaining in CON_ and MUT_*BRCA2* iPSCs. (**C-D**) Relative gene expression profiles (qPCR) of iPSC-specific genes; *OCT4* and *SOX2* in CON_ and MUT_*BRCA2* iPSCs. (**E-F**) Relative gene expression profiles (qPCR) of both *BRCA2* and *BRCA1* in CON_ and MUT_*BRCA2* iPSCs. (**G-H**) Ki67 positive proliferating nuclei analysis in CON_ and MUT_*BRCA2* iPSCs, representative images, and quantitation. (**I**) Represented IF images showing the F-actin organization, localization of apical (CK8/18), and basal (p63) polarity markers in CON_ and MUT_*BRCA2* iPROS. Scale bars: 100 mM (A, F) and 40 mM (J). (n = 3 different biological replicates). Significance calculated with pairwise comparison (t-test; Welch correction); ns= p>0.05, *p < 0.05.

**Figure S3:**
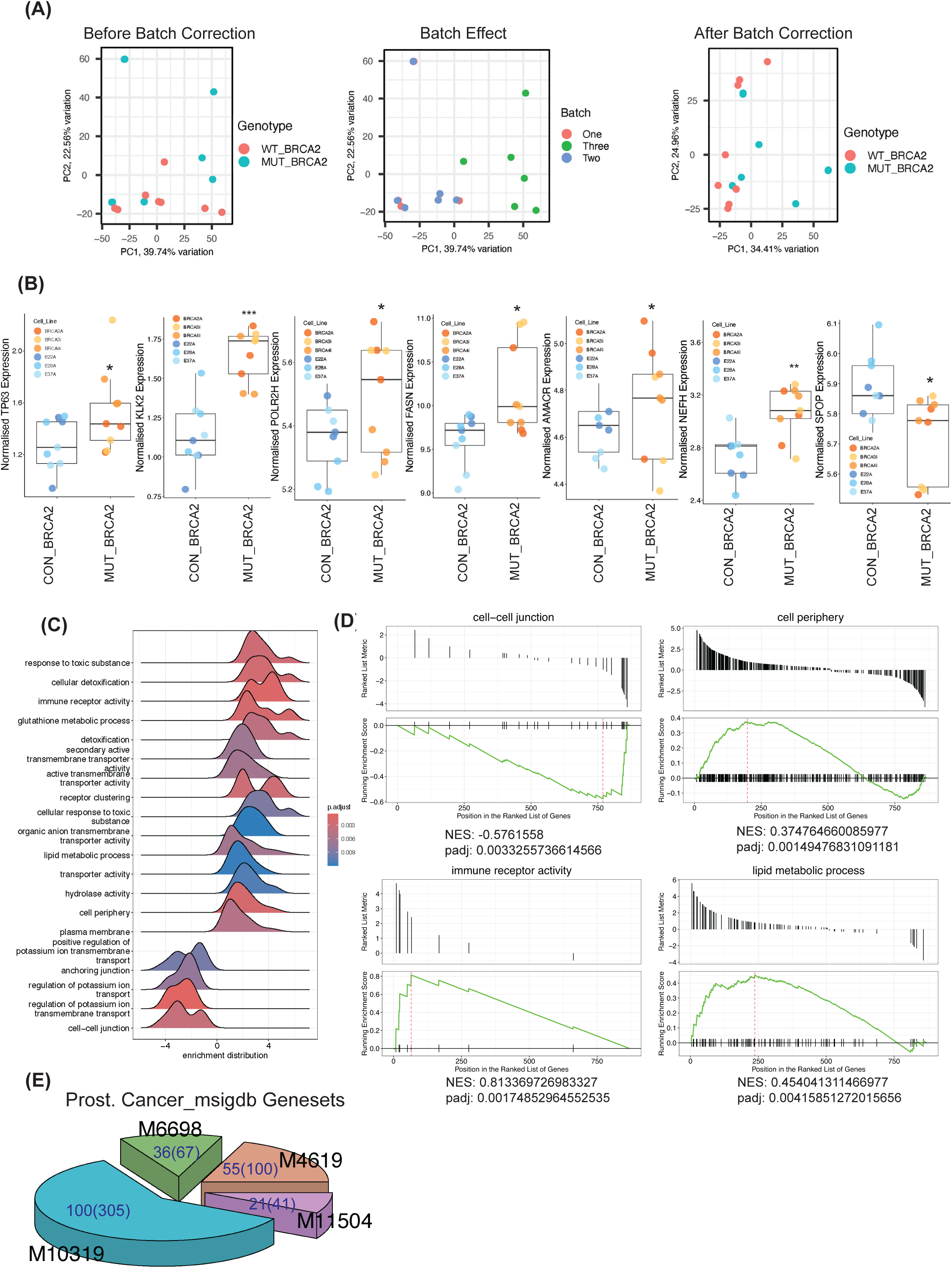
(**A**) PCA analysis and batch correction showing the variance among samples (explained in the result). (**B**) Normalized expression of prostate- and PCa-specific genes. (**C**) GSEA ridge map with significant DEGs between CON_ and MUT_*BRCA2* iPROS samples. (**D**) GSEA barcode plots with significant DEGs between CON_ and MUT_*BRCA2* iPROS samples. (**E**) Pie chart showing gene commonality/overlapping in MUT_BRCA2 with four different MSigdb PCa datasets on significant DEGs between MUT_ vs. CON_BRCA2 iPROS samples.

**Figure S4:**
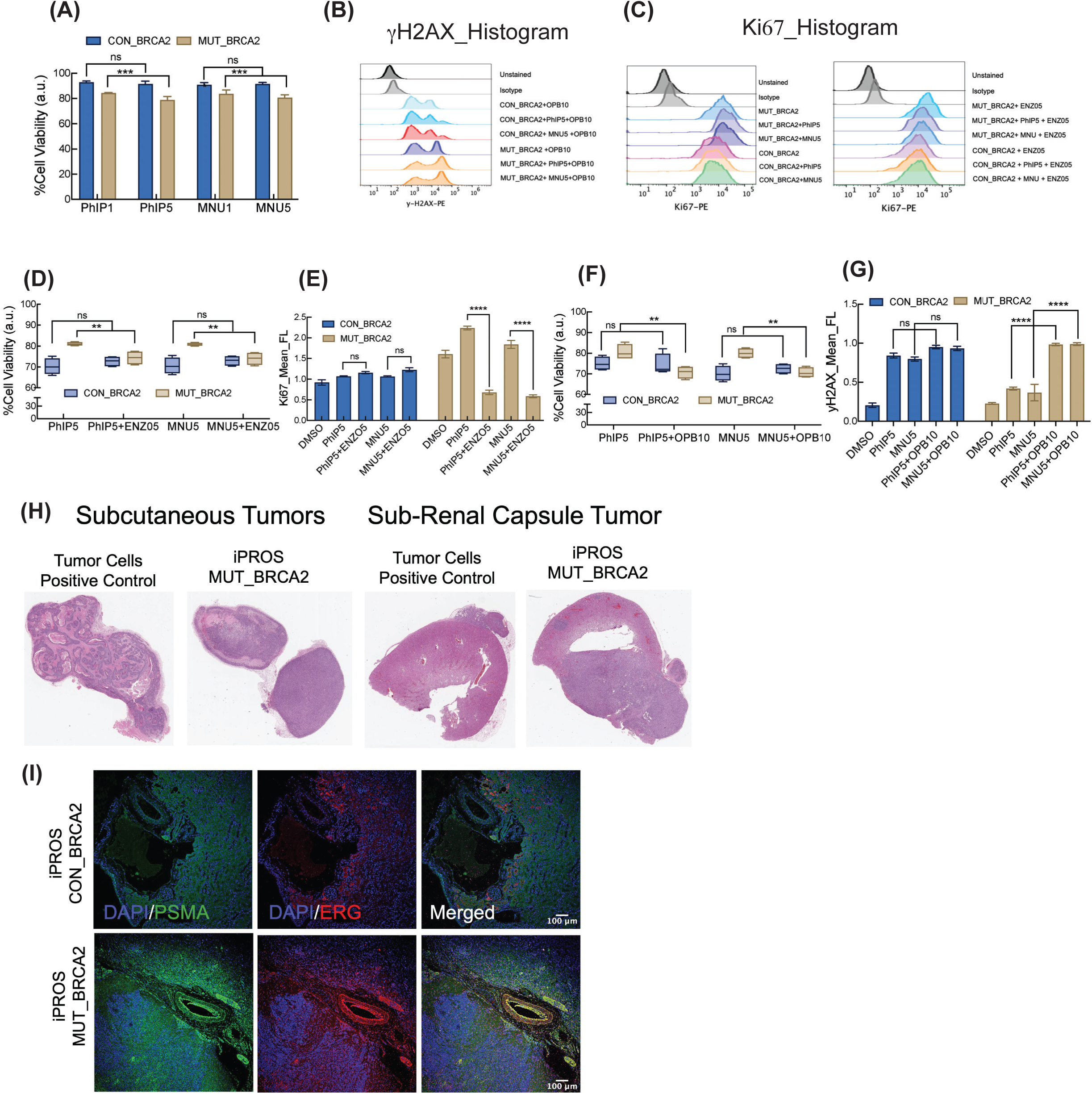
(**A**) Bar-plot showing the cytotoxicity of CON_ and MUT_BRCA2 iPROS after 3 weeks of incubations with PhIP and MNU. (**B-C**) Histograms showing the flow cytometric analysis of γH2AX and Ki67 in PhIP and MNU treated CON_ and MUT_BRCA2 iPROS. (**D-E**) Boxplots showing the cytotoxicity and Bar-plot showing the proliferation of CON_ and MUT_BRCA2 iPROS at 4^th^ week following 2 weeks after PhIP and MNU treatment withdrawal and addition of Enzalutamide for 2wks., by LDH cytotoxic assay, and mean-fluorescence obtained by flow cytometry. (**F**) LDH cytotoxic assay showing the cytotoxicity of CON_ and MUT_BRCA2 iPROS at 4^th^ week following 2 weeks after PhIP and MNU treatment withdrawal and addition of Olaparib for 2 weeks. (**G**) Flow cytometric analysis of γH2AX for detecting DNA damage in iPROS with and without Olaparib treatment at 4^th^ week. (**H**) Representative H&E staining of whole mount of subcutaneous and renal capsule tumor sections obtained from positive control and MUT_BRCA2 iPROS. Boxed sections are separately shown in Fig.4K (**I**) Representative IF staining of whole mount of kidney with PSMA and ERG, counterstained with DAPI. Significance was calculated using 2-way ANOVA with multiple comparisons: ns= p>0.05, *p < 0.05, **p < 0.01, ***p < 0.001, ****p < 0.0001.

